# Increased mGlu5 mRNA expression in BLA glutamate neurons facilitates resilience to the long-term effects of a single predator scent stress exposure

**DOI:** 10.1101/2020.09.23.310342

**Authors:** John Shallcross, Lizhen Wu, Lori A. Knackstedt, Marek Schwendt

## Abstract

Post-traumatic stress disorder (PTSD) develops in a subset of individuals exposed to a trauma with core features being increased anxiety, and impaired fear extinction. To model the heterogeneity of PSTD behavioral responses, we exposed Sprague-Dawley rats to predator scent stress (TMT) once for 10 minutes and then tested for anxiety-like behavior 7 days later using the elevated plus-maze and acoustic startle response. Rats displaying anxiety-like behavior in both tasks were classified as stress-Susceptible, and rats exhibiting behavior no different from unstressed Controls were classified as stress-Resilient. Our previous findings revealed increased mRNA expression of mGlu5 in the amygdala and PFC and CB1R mRNA in the amygdala of Resilient rats. Here, we performed fluorescent in situ hybridization (FISH) to determine the subregion and cell-type-specific expression of these genes in Resilient rats. We found higher mRNA expression of mGlu5 in the BLA, IL, and PL, and CB1R in the BLA of Resilient rats relative to Controls. Using dual-labeled FISH we determined that mGlu5 and CB1R mRNA increases were limited to vGlut+ cells. To test the necessity of mGlu5 receptor activity for attenuating contextual fear, intra-BLA infusions of the mGlu5 negative allosteric modulator MTEP were administered prior to context re-exposure. MTEP increased contextual fear on the day of administration, which extinguished over the course of two additional un-drugged sessions. These results suggest that an enhanced mGlu5 expression within BLA glutamate neurons contributes to the behavioral flexibility observed in stress-Resilient animals by facilitating a capacity for extinguishing contextual fear associations.

## Introduction

Approximately 8.3-25% of trauma-exposed adults develop a symptom profile meeting full DSM-5 diagnostic criteria for post-traumatic stress disorder (PTSD), reflecting interindividual functional alterations within fear regulating neural systems (Breslau et al. 1991; Perkonigg et al. 2000; Sareen 2014). A central underlying feature of PTSD is the failure to reduce reactivity to trauma-conditioned stimuli following repeated presentations in the absence of actual threat, also characterized as fear extinction (Orr et al. 2000; Guthrie and Bryant 2006; Bouton et al. 2006; Blechert et al. 2007). Considerable evidence from both human and rodent studies implicates the coordinated activity of the amygdala and medial prefrontal cortex (mPFC) in fear extinction (Sotres-Bayon et al. 2006; Koenigs and Grafman 2009; Bukalo et al. 2015). The amygdala comprises distinct subnuclei, including the central nucleus (CeA) and the basal lateral nucleus (BLA) (Duvarci and Pare 2014). The CeA contains mainly GABAergic medium spiny neurons and serves to mediate the autonomic stress response via direct descending projections to hypothalamic and brainstem nuclei (Sah et al. 2003; Ciocchi et al. 2010). The BLA is comprised of glutamatergic primary neurons (PN) and GABAergic interneurons (IN), and supports the acquisition and maintenance of extinction memory (Gale et al. 2004; Ehrlich et al. 2009; Pape and Pare 2010). The mPFC contains both PN and IN cell types and can be subdivided into prelimbic (PL) and infralimbic (IL) regions which function dichotomously to promote the expression and extinction (respectively) of learned fear (Sierra-Mercado et al. 2011; McGarry and Carter 2017; Caballero et al. 2019). Current models indicate that consolidation of extinction memory relies on BLA ↔ IL interactions driven through direct reciprocal PN projections (Herry et al. 2008; Laricchiuta et al. 2016; Bloodgood et al. 2018). Moreover, extinction-evoked bidirectional changes in excitability across BLA ↔ IL circuits contribute to the suppression CS-evoked fear response (Day et al. 2004; Vouimba and Maroun 2011; Kheirbek et al. 2013; Cho et al. 2013; Pietrzak et al. 2014; Kim and Cho 2017).

Recent evidence from clinical studies suggests that the prevalence of PTSD-related symptoms may be linked to the availability of mGlu5 (Holmes et al. 2017; Esterlis et al. 2018) and CB1 (Neumeister et al. 2013; Pietrzak et al. 2014) receptors. Generally located on postsynaptic membranes at peri-synaptic and synaptic sites (Rodrigues et al. 2002; Muly et al. 2003; Fitzgerald et al. 2019), the Gα_q/11_-coupled mGlu5 receptors exert control over intrinsic neuronal excitability and structural synaptic plasticity (Heuss et al. 1999; Ayala et al. 2009; Wang and Zhuo 2012; Chen et al. 2017). Moreover, studies in rodents demonstrate that activation of mGlu5 receptors in the BLA (Rudy and Matus-Amat 2009; Mao et al. 2013; Rahman et al. 2017) and mPFC (Fontanez-Nuin et al. 2011; Sepulveda-Orengo et al. 2013) is necessary for ‘normal’ fear extinction behavior. Using a rodent model of PTSD involving predator scent stress exposure (PSS), we have found that rats resilient to the long-term anxiety and conditioned fear produced by a single brief TMT (fox pheromone) exposure display increased mGlu5 mRNA expression in the mPFC and amygdala relative to both unstressed controls and rats susceptible to anxiety and conditioned fear (Schwendt et al. 2018). Such increases are observed 24 days after TMT exposure when rats are killed 2 hours after assessment of conditioned contextual fear (freezing in the PSS environment). Accompanying such increases in mGlu5 mRNA expression are reduced contextual fear relative to the stress-susceptible rats that exhibit robust contextual fear.

Like mGlu5, the Gα_i/o_-coupled type 1 cannabinoid receptor (CB1R) regulates neuronal excitability and synaptic plasticity and has been shown to play a crucial role in fear extinction (Marsicano and Lutz 1999; Kano et al. 2009; Vogel et al. 2016; Krabbe et al. 2018; Zou and Kumar 2018; Shen et al. 2019). In the BLA, CB1 receptor mRNA is highly expressed in a sparsely distributed population of cholecystokinin positive (CCK+) GABAergic interneurons that synapse locally onto projection PNs (Katona et al. 2001; Jasnow et al. 2009; Yoshida et al. 2011). At these sites, reciprocal functional interactions between presynaptic CB1 and postsynaptic mGlu5 receptors produces transient disinhibition of PNs which serves to maintain tight regulatory control over the excitability of bidirectional PL and IL targeting BLA outputs (Zhu and Lovinger 2005; Vogel et al. 2016; Marcus et al. 2020). In the PSS model, we have found increased CB1 receptor expression in the amygdala of stress-resilient rats relative to both unstressed controls and stress-susceptible rats (Schwendt et al. 2018).

In addition to finding increased mGlu5 and CB1 mRNA expression in stress-Resilient rats, our prior work finds that Susceptible but not Resilient outbred Sprague-Dawley rats display contextual fear that does not extinguish (Schwendt et al. 2018; Shallcross et al. 2019) We also found that systemic administration of the mGlu5 positive allosteric modulator CDPPB reduces contextual fear, but not anxiety, in Susceptible rats, further supporting a role for mGlu5 in mediating PSS conditioned fear (Shallcross et al. 2019). The current study seeks to further explore the potential role for mGlu5 expression in resistance to contextual fear. Here, we utilize fluorescent in situ hybridization (FISH) to examine sub-regional and cell-type-specific mGlu5 mRNA expression in the CeA, BLA, PL, and IL of rats resilient to the long-term effects of predator scent stress. In addition to mGlu5, we measured mRNA expression of functionally related CB1R in BLA PN and IN cell populations. Finally, we show that pharmacological inhibition of mGlu5 receptors in the BLA increases fear responding in Resilient rats during re-exposure to the predator scent stress context.

## Materials and Methods

### Animals

Procedures were approved by the Institutional Animal Care and Use Committee at the University of Florida. Adult male Sprague-Dawley rats (Charles River Laboratories, Wilmington, MA, USA) were single-housed in a ventilated rack system and maintained on a reverse 12:12 light-dark cycle (lights off at 7:00 am) with mild food restriction (20 g/day). Rats were carefully handled by both male and female experimenters during a one-week acclimation period. Experiments were performed at the beginning of the dark phase. Forty-two rats were used for these experiments.

### Predator-Scent Stress Exposure

Stressed rats were exposed to 12,3,5-Trimethyl-3-thiazoline (TMT; BioSRQ, Sarasota, FL, USA), a fox pheromone, or the control unscented condition. On Day 1, TMT (3 μl) was pipetted onto filter paper positioned underneath the stainless-steel mesh flooring of a clear cylindrical plexiglass chamber (35 cm height x 20 cm radius). Rats were individually placed in the center of the chamber for a 10-minute exposure. Rats were not in physical contact with the TMT. Rats were then removed and returned to the home cage.

### Tests for anxiety-like behavior

Seven days after TMT exposure, rats were tested on the elevated plus maze (EPM) and acoustic startle response (ASR) task to evaluate anxiety-like behavior. The EPM apparatus (Med Associates, St. Albans, VT, USA) consists of four black plexiglass arms (50 cm length x 10 cm width) raised 72 cm from the floor. Two open arms (1.5 cm high walls) and two closed arms (50 cm high walls) are joined by a center square platform (10 cm x 10 cm) illuminated at 50 lux. Rats were individually placed on the center platform and given 5 min to explore the maze. Time spent in the open arms (OA) was measured using EthoVision XT 14 software (Noldus Information Technology, Leesburg, VA). Within 20 min of EPM testing, the acoustic startle response (ASR) was assessed. Four ventilated soundproof chambers (San Diego Instruments, San Diego, CA) contained a transparent plexiglass cylinder resting on a pressure-sensitive platform. The maximum response amplitude was registered during presentation of acoustic stimuli. The startle habituation program consisted of 5 min acclimation followed by thirty 110 dB white noise pulses (ITI: 30 – 45 sec). Following EPM/ASR tests, rats were phenotyped as stress-Susceptible, Resilient, or Intermediate (see below and (Schwendt et al. 2018; Shallcross et al. 2019). Only Control rats and TMT-exposed rats meeting the criteria for stress-Resilience were assigned to Experiment 1 or Experiment 2.

### Experiment 1: Determination of amygdala and mPFC subregion and cell-type specificity of mGlu5 mRNA upregulation in Resilient rats

On Day 21 (14 days after EPM/ASR testing; Fig. 1A), Control rats (n=11) and a subset of Resilient rats (n=10) were re-exposed to the TMT-conditioned chamber in the absence of TMT for a 10-minute contextual fear test. A second subset of Resilient rats (n=6) were instead placed into a neutral environment. In accordance with the timing of our previous publication (Schwendt et al. 2018), 2 hr later, rats were administered an overdose of sodium pentobarbital solution (100 mg/kg IP). Transcardial perfusions were performed using nuclease free 0.9% NaCl followed by cold 4% paraformaldehyde (PFA) in PBS. Brains were post-fixed for 12 hr at 4°C in 4% PFA in PBS, cryopreserved in 30% sucrose, and stored at −80°C after 24-48 hrs. Coronal brain sections (12 μm) containing the BLA (−2.04 to −2.76 mm relative to Bregma) and mPFC (−2.04 to −2.76 mm relative to Bregma, according to (Paxinos and Watson 2007, Fig. 1B) were obtained using a freezing cryostat (Leica CM1950). Tissue sections were rinsed in PBS, mounted onto positively charged glass slides (Superfrost Plus Gold), air-dried for 20 min at RT, and stored at −80°C. Tissue from all Resilient-Neutral rats was used for both amygdala and PFC regions. However, due to technical problems hindering slicing of the dmPFC from the first cohort of rats, separate cohorts of Control and Resilient re-exposed rats were used for BLA and mPFC (Control BLA n=5; Control mPFC n=6; Resilient re-exposed BLA n=5; Resilient re-exposed mPFC n=5).

**Figure 1.**
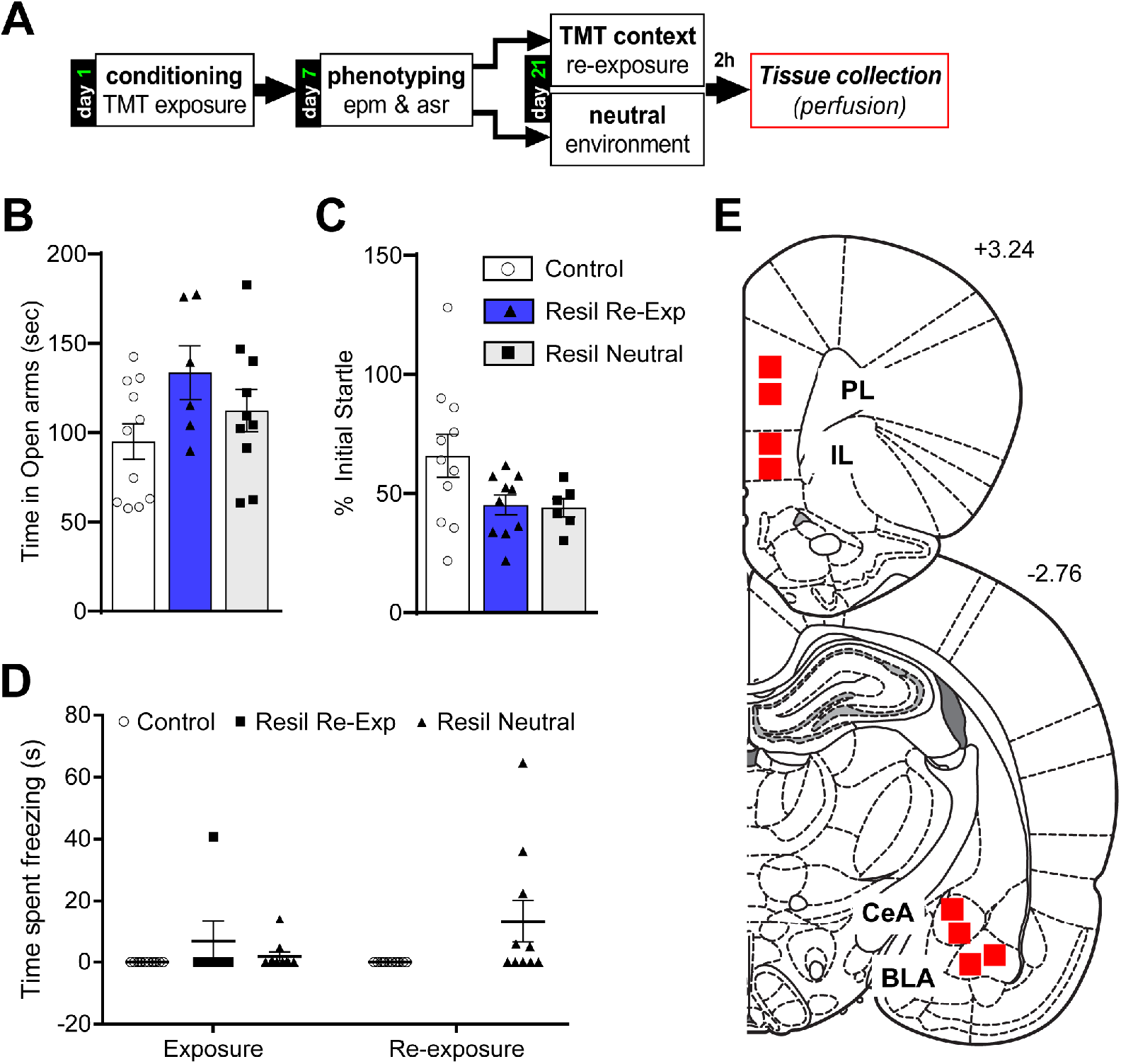
Experimental design and behavior for Experiment 1. **A.** Timeline of Experiment 1. **B.** Time spent in the Open Arms (OA) of the elevated plus maze (EPM) did not differ between groups. **C.** Habituation of the acoustic startle response (ASR) did not differ between groups. **D** A main effect of group [Control vs. Resilient (Resil)] was observed for time spent freezing during TMT exposure and re-exposure to the TMT context. **E.** Individual sub-regions selected for the FISH analysis in the amygdala and prefrontal cortex for Experiment 1, as highlighted on rat brain coronal sections (according to: Paxinos and Watson, 2007).

#### Fluorescent in situ Hybridization (FISH)

Fluorescent in situ hybridization of mRNA was performed using the RNAscope Multiplex Fluorescent Reagent Kit (Advanced Cell Diagnostics; Newark, CA, USA) according to the manufacturer’s instructions with some modifications (Wang et al. 2012). Briefly, slide-mounted sections were warmed to 60°C, fixed in cold 4% PFA in PBS (90 min), and dehydrated using ethanol gradient. Slides were treated with target retrieval reagent (Advanced Cell Diagnostics) followed by proteinase digestion with pretreatment #3 for 30 min under humidity-controlled conditions. mRNA probe hybridization and signal amplification was performed for mGlu5 and CB1 alone and in combination with both vGlut1 and GAD65. The following probes (Advanced Cell Diagnostics) were used: mGlu5 (GRM5; Cat. #471241), CB1 (CNR1; Cat. #412501), vGlut1 (SLC17A7; Cat. #317001), and GAD65 (GAD2; Cat. #43580). Amplifier 4 Alt-A was selected to fluorescently label mGlu5 and CB1 probes with ATTO 550 (orange) dye and vGlut1 and GAD65 with Alexa 488 (green). Cell nuclei were counterstained with DAPI (4’,6-diamidino-2-phenylindole), and antifade mounting media (ProLong Gold) was applied with coverslip.

#### Image acquisition and analysis

Fluorescent confocal images were obtained using an Olympus DSU (DU-DBIX) Spinning Disk Confocal Module mounted on an Olympus IX81 Inverted Fluorescent Microscope Frame fitted with a Hamamatsu ORCA-Flash 4.0LT+ (Model # C11440-42U30) monochrome 16-bit camera. A CoolLED Model # pE-300white fluorescent lamp was used in combination with Semrock excitation filters (DAPI, FITC, & TRITC) for visualization of mRNA probe fluorescence. Positive and negative control slides were used to calibrate baseline exposure LED intensity values. DAPI and FISH probes were sequentially scanned. Image stacks 10 μm in depth were acquired at 1 μm intervals using an Olympus 40x UPLSAPO dry objective and Olympus cellSens Dimension v2 Software. For each rat, two non-overlapping regions of interest (ROI) were captured from each of the three brain regions (BLA, IL, PL) from two separate slices. Stacks were then loaded into ImageJ (Schneider et al. 2012), and co-localization of target mRNA and cell type marker mRNA was manually performed using the ImageJ “Multi-point” tool. Cells were identified based on DAPI stain, and FITC/TRITC channels were toggled on or off to assess overlapping fluorescence in the same cells across the Z-axis. Detection of neuronal type marker mRNA signal on at least three focal Z-planes was required for within-cell mRNA expression analysis. Qualifying cells were numbered on a maximum intensity Z-projection from each ROI stack, which served as a reference for quantification analysis with TransQuant software (Bahar Halpern and Itzkovitz 2016). TransQuant is a GUI based software for Matlab that counts fluorescent puncta representative of single-molecule mRNAs across three-dimensional image stacks. Total expression of Cy3-labeled target mRNAs (mGlu5, CB1R) was quantified for each ROI image stack, and two separate stacks (ROIs) were analyzed from each brain section for every rat. The average mRNA expression across two representative brain slices from each rat was calculated. For cell-type-specific within-cell mRNA expression analysis, reference images were used to identify and manually segment each cell co-expressing both FITC- and Cy3-labeled mRNA within each ROI. The total number of Cy3 labeled mRNA per individual cell was counted and averaged across all cells for each brain area.

### Experiment 2: Testing the functional relevance of BLA mGlu5 receptors in mediating conditioned fear extinction

On the day following EPM/ASR testing (Fig. 4A), 15 Resilient rats were anesthetized with ketamine HCl (87.5 mg/kg, IP) and xylazine (5 mg/kg, IP) and bilateral guide cannulae (20 gauge, Plastics One) targeting the BLA (AP: −3.3 mm, ML: ±4.9 mm, DV: −6.3 mm, all relative to Bregma (Paxinos and Watson 2007) were implanted and secured to the skull with screws and dental cement. The coordinates for cannula placement were determined from the rat brain atlas (Sareen 2014). Rats were administered Carprofen (2.5 mg/kg) for analgesia on the day of surgery and for two days following.

On Day 21 (14 days after EPM/ASR; Fig. 4A), rats underwent once daily contextual extinction trials for three consecutive days. A syringe pump (Harvard Apparatus) was connected to Hamilton syringes attached to 33-gauge injectors Thirty minutes before testing microinjectors were inserted into cannulae, extending 2 mm below the cannulae, and either vehicle or 3-((2-Methyl-4-thiazolyl)ethynyl) pyridine (MTEP) hydrochloride (Abcam Biochemicals, Cambridge, MA, USA) were infused over 2 min. The vehicle for MTEP was 1% Tween-80 in artificial cerebrospinal fluid (in mM: 125 NaCl, 1 CaCl_2_, 2.5 KCl, 4 MgCl_2_, 25 NaHCO3, 1.25 NaH_2_PO_4_, 0.4 ascorbic acid and 10 D-glucose, pH 7.2–7.4). MTEP (10 μg/side) or vehicle was delivered into the BLA at a rate of 0.5 μl/min in a total volume of 0.5 μl/side. This concentration was chosen based on a previous study that found intracranial infusions targeting the amygdala with 10 μg/side of MTEP impaired extinction memory (Mao et al. 2013).

Rats were then returned to the TMT exposure chambers (in the absence of TMT) for 10 min contextual fear trial (Extinction Day 1). On Extinction Day 2 and 3, all rats received vehicle injections prior to placement into the chamber. Immediately following the Day 3 trial, transcardial perfusions were performed using 0.9% saline and 4% and brains were extracted, post-fixed, sectioned (30 μm), and stained with Cresyl violet dye to verify cannulae placement under a light microscope.

### Data analysis

Startle habituation was calculated as the percent change in startle amplitude from the average of the first six trials to the average of the last six trials as in our previous publications (Schwendt et al. 2018; Shallcross et al. 2019). A double median split was conducted on EPM (time spent in the OA) and ASR (% habituation of initial startle). GraphPad Prism (version 6.0) was used for statistical analysis with the alpha level set at p ≤ 0.05. One-way ANOVA and unpaired t-tests were used to compare phenotypic differences in EPM and ASR behavior for rats used in Experiment 1-2 and for within region comparison of mRNA expression between treatment groups. Freezing in the TMT context was compared between Resilient and Control rats with two-way mixed factorial repeated-measures (RM) ANOVAs with Group as the between-subjects factor and Time as a within-subjects factor. Significant interactions were followed by Tukey’s post hoc analyses with corrections for multiple comparisons.

## Results

### TMT exposure induces variable long-term anxiety response

The Resilient rats used here were part of a cohort of 307 rats exposed to TMT and tested in EPM/ASR 7 days later, as described above (Fig 1A). The median time spent in the OA of the EPM was 56.2 sec and the median habituation of the ASR was 61.2%. Rats that scored below the median time spent in the OA and above the median ASR habituation were phenotyped as Susceptible; those scoring above-median OA time and below ASR habituation were phenotyped as Resilient (Resil) as in our previous report (Schwendt et al. 2018). All Susceptible and some Resilient rats were used in a previous publication (Shallcross et al., 2019) whereas a subset of Resilient rats was used for the present set of experiments (n=31) along with a new cohort of unstressed Control rats (n=11).

### Experiment 1

#### Anxiety-like and fear behavior

EPM and ASR data from the entire population of TMT-exposed rats have been published previously (Shallcross et al., 2019). There were no differences in time spent in the OA or percent habituation in the ASR between Resilient and Control rats used in the present experiment (Fig. 1BC). A two-way ANOVA compared time spent freezing on the day of TMT/Control exposure and the day of re-exposure, finding a main effect of Group [F_(1, 20)_ = 5.621, p=0.0279; Fig. 1D], with TMT-exposed Resilient rats displaying greater freezing than Controls. Data from Resilient rats not re-exposed to the TMT-context are displayed in Fig. 1D, but were not analyzed in the ANOVA as there was no data for a re-exposure to be analyzed.

#### Resilient rats display increased mGlu5 mRNA levels in the BLA, PL, and IL

In a previous report, we revealed elevated mGlu5 mRNA in mPFC, and amygdala tissue punches taken from resilient rats following a contextual fear test. Given subregion-specific differential signaling dynamics in both mPFC and amygdala, we sought to expand on these findings using single-molecule fluorescence in situ hybridization (FISH) for labeling of mGlu5 mRNA in mPFC and amygdala brain slices from control rats and resilient rats re-exposed to either fear or neutral associated context on experimental Day 21 (see Fig. 1A for timeline). Improved spatial resolution with FISH permitted detection of mRNA puncta within subregions of interest (Fig. 1E, 3D). Robust uniform distribution of fluorescent puncta was observed in PL and IL regions in both neutral and fear context exposed resilient rats (Fig. 2B). Even distribution of mGlu5 fluorescence was also observed in the CeA and the BLA, however, mGlu5 signal in the BLA was markedly more intense (Fig. 2B).

**Figure 2.**
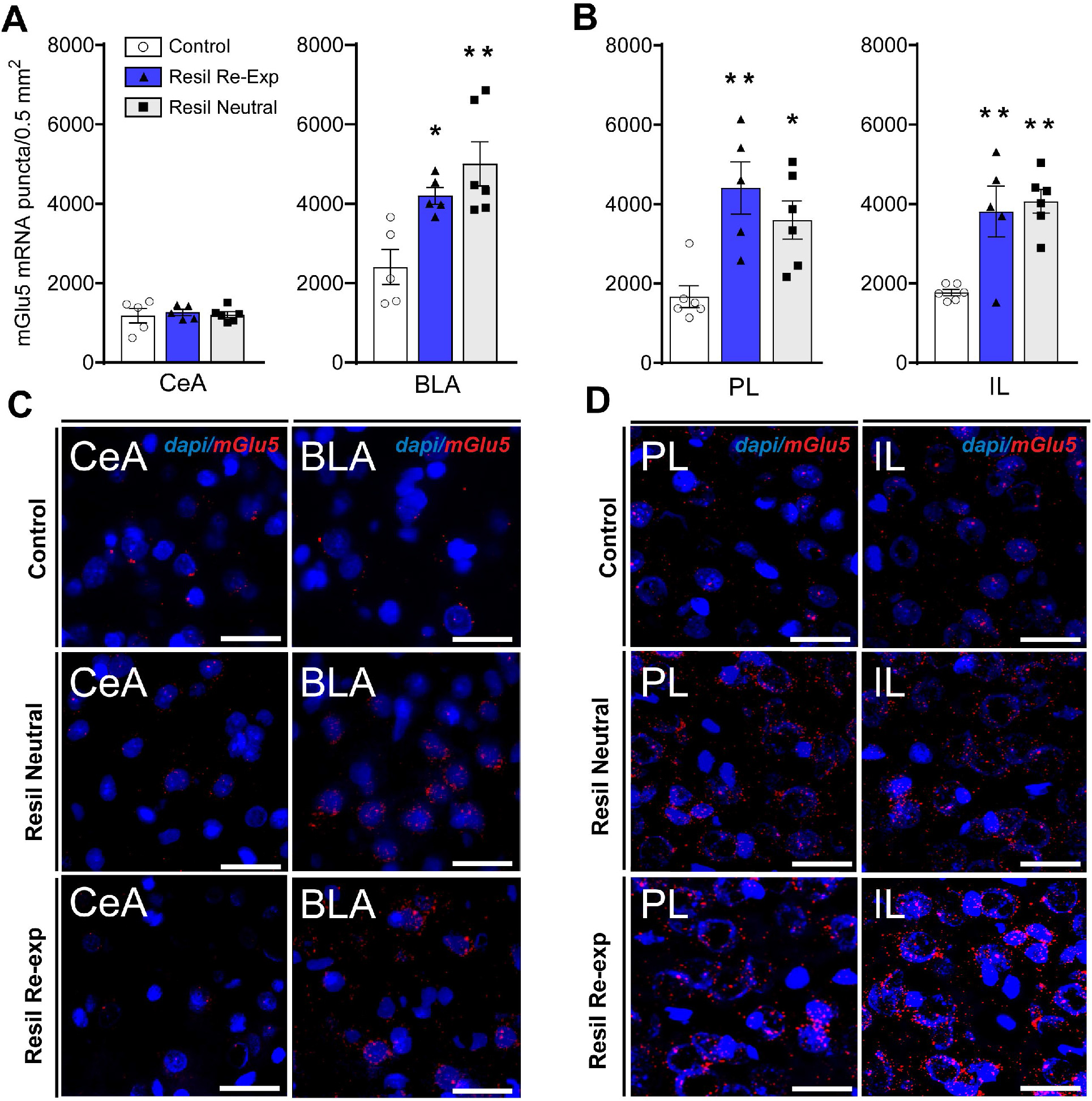
Resilience to the long-term effects of a single predator scent stress exposure is accompanied by increased mGlu5 mRNA expression in several brain regions. **A.** mGlu5 mRNA expression was greater in Resilient (Resil) rats (regardless of re-exposure to the TMT context) in the BLA but not the CeA. **B.** mGlu5 mRNA expression was greater in Resilient rats (regardless of re-exposure to the TMT context) in both the PL and IL regions of the mPFC. **C.** Representative images from the CeA and BLA. **D.** Representative images from the PL and IL. * = p<0.05 compared to Control; ** = p<0.01 compared to Control. Scale bar = 40 μm.

There were no group differences in mGlu5 mRNA expression in the CeA (Fig. 2A,C left). A main effect of group was found for mGlu5 expression in the BLA [F_(2, 13)_ = 8.714, p = 0.004; Fig 2A,C right]. Post-hoc analysis revealed both neutral (p<0.01) and TMT-context exposed (p<0.05) displayed greater mGlu5 expression than Controls. Quantification of puncta indicated a main effect of Group for mGlu5 mRNA expression in both PL [F_(2, 14)_ = 8.766, p=0.0034] and IL [F_(2, 14)_ = 11.84, p=0.001]. Post hoc analyses revealed increased mGlu5 mRNA levels in both neutral (PL, p < 0.05; IL p < 0.01] and context re-exposed (PL, p < 0.01; IL, p < 0.01) Resilient rats relative to controls (fig. 1e). Exposure to the fear associated context did not result in different levels of mGLu5 mRNA relative to neutral context exposure in BLA, PL or IL.

#### Increased levels of mGlu5 mRNA in Resilient rats is restricted to glutamate neurons

The mGlu5 receptor is expressed in several cell types throughout the CNS, including both glutamatergic and GABAergic neurons (Ferraguti and Shigemoto 2006; Fitzgerald et al. 2019). To determine in which cell subtype mGlu5 expression is changed in Resilient rats, co-localization of mGlu5 mRNA with the GABA synthesizing enzyme GAD65 (GAD2) and vesicular glutamate transporter coding vGlut1 (SLC17A7) was examined in the BLA, PL, and IL of Resilient rats and Controls. The CeA was not examined, as no group differences in overall mGlu5 expression were observed. Consistent with previous reports, we observed vGlut1 labeling in the majority of BLA cells (vGlut1 +, Fig. 3C), with sparse distribution of cells reporting GAD65 (GAD65+, not shown). Almost all vGlut1 + (98%) and GAD65+ (95%) cells co-localized with mGlu5 mRNA. In 14-21% of mGlu5+ cells, co-localization with GAD65 was observed, while the remaining mGlu5+ cells contained vGlut1 mRNA. Using TransQuant, we segmented vGlut1+ and GAD65+ cells and quantified mGlu5 puncta within cytoplasmic boundaries using DAPI nuclear counterstain for reference (see Fig. 2C Resilient-Neutral IL, for example). mGlu5 expression within vGlut1+ cells differed between groups in the BLA [F_(2, 13)_ = 7.452, p = 0.007]; PL [F_(2, 14)_ = 20.27, p < 0.001]; and IL [F_(2, 17)_ = 5.582, p=.014]. Post hoc analyses revealed elevated mGlu5 mRNA in vGlut1+ cells across all three regions in both neutral (BLA p < 0.05; PL p < 0.01; IL p < 0.01) and re-exposed (BLA p < 0.05; PL p < 0.01; IL p < 0.05) Resilient rats relative to Control. The amount of mGlu5 in vGlut1+ cells did not differ between fear or neutral context re-exposed rats in any brain region. Quantification of mGlu5 puncta within GAD65+ cells revealed no group differences in any region analyzed.

**Figure 3.**
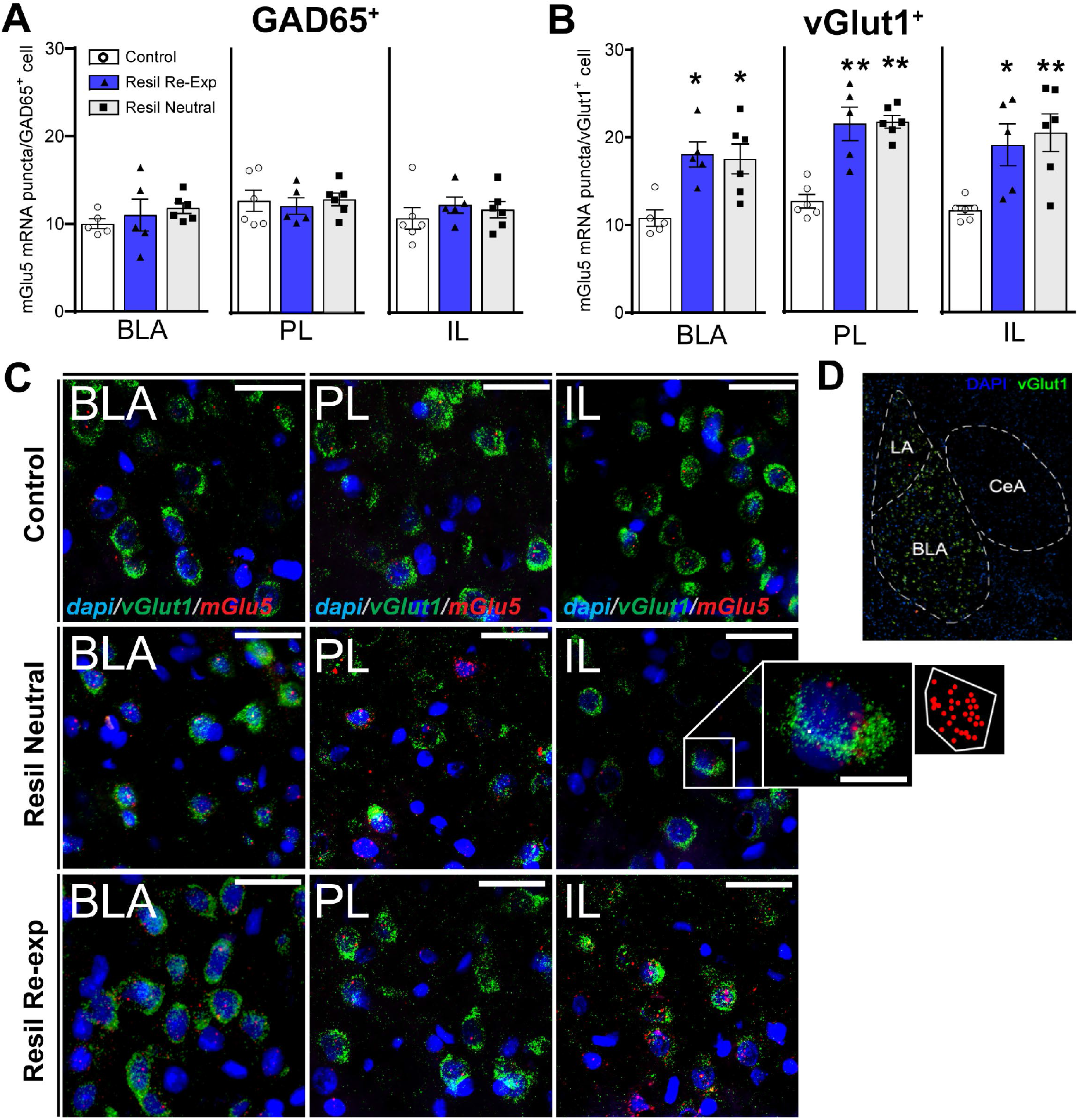
Resilience to the long-term effects of a single predator scent stress exposure is accompanied by increased mGlu5 mRNA expression in glutamatergic but not GABAergic neurons. **A.** mGlu5 mRNA expression in GAD67+ cells did not differ between groups. **B.** Relative to Control rats, mGlu5 expression was greater in vGlut1+ cells in both Resilient (Resil) rats that were re-exposed to the stress context and those that were not. **C.** Representative images from BLA, PL, and IL of each group. **D.** Delineation of amygdala regions. * = p<0.05 compared to Control; ** = p<0.01 compared to Control. Scale bar = 40 μm.

#### Elevated CB1 mRNA in resilient rats is co-localized with vGlut1

We have also previously reported elevated amygdala CB1 receptor (CB1R) mRNA expression in the amygdala of fear context re-exposed Resilient rats. Here we conducted FISH for CB1R (CBR1) mRNA in amygdala slices as done for mGlu5. Distribution of CB1R mRNA in the BLA was consistent with previous reports describing a large population of low CB1 mRNA expressing cells, and a sparse population of high CB1 expressing cells (Fig. 4B) Total level of CB1 puncta in the BLA differed between groups [F_(2, 13)_ = 21.56, p<.001]. Post hoc analysis revealed higher CB1 expression in both re-exposed (p < 0.001) and neutral (p < 0.001) Resilient rats relative to controls (Fig. 4A left). Since CB1 mRNA is expressed in both excitatory and inhibitory BLA cell subtypes, we assessed CB1 in vGlut1+ and GAD65+ neurons as we did for mGlu5. We found that most (90-95%) CB1 high-expressing cells co-localized with GAD65+ (although not all GAD65+ co-localized with CB1), and most (92-95%) CB1 low-expressing cells co-localized with vGlut1+ (although not all vGlut1+ co-localized with CB1) (Fig. 4B). Segmentation of cells and quantification of CB1 puncta within cell subtypes indicated a main effect of Group on CB1 expression in vGlut1+ cells [F_(2, 13)_ = 9.387, p < 0.001] Post hoc analysis revealed that both neutral (p < 0.01) and fear (p < 0.01) context exposed resilient rats expressed higher CB1 puncta in vGlut1+ (low expressing) cells relative to controls (Fig. 4B right). Quantification of CB1 puncta within GAD65+ (high expressing) cells revealed no group differences (Fig. 4B center).

**Figure 4.**
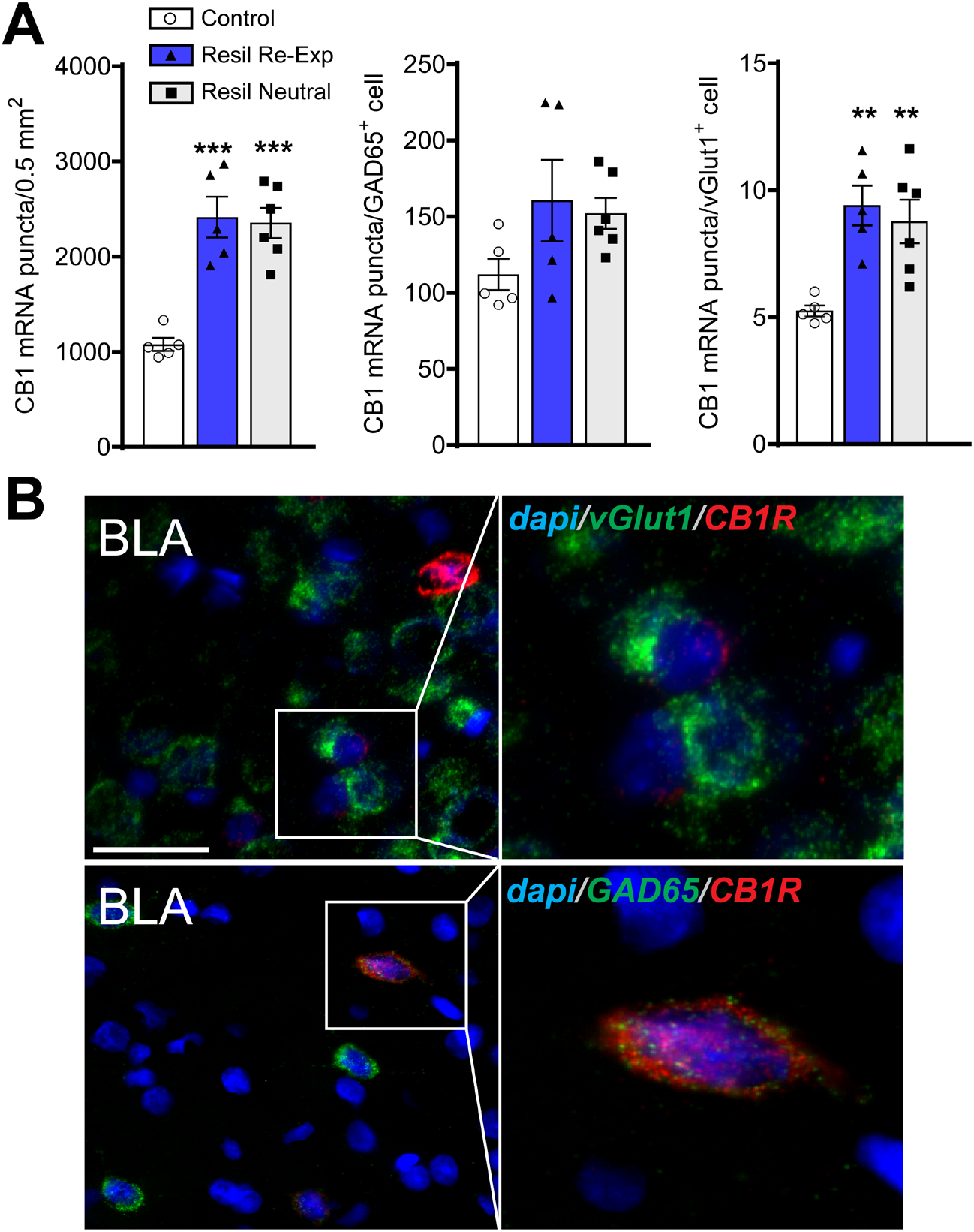
CB1 mRNA expression is upregulated in glutamatergic but not GABAergic BLA neurons in stress-Resilient rats. **A.** Resilient (Resil) rats show upregulation of CB1 mRNA in the BLA overall (left), and in vGlut1+ cells (right), but not in GAD65+ cells (center). **B.** Representative images of CB1 mRNA in combination with vGlut1 (top) and GAD65 (bottom). ** = p<0.01 compared to Control; *** = p<0.001 compared to Control. Scale bar = 40 μm.

#### mGlu5 inhibition increases context conditioned fear expression in resilient rats

We tested the hypothesis that mGlu5 receptors mediate inhibition of conditioned fear displayed by Resilient rats during fear context re-exposure (see Fig. 5A for timeline). One (MTEP) rat was eliminated for cannulae placement outside the BLA and one (Vehicle) rat was eliminated for displaying freezing behavior on all three days that was more than 2 standard deviations from the mean, leaving 7 rats in the MTEP group and 6 rats in the Control group. There were no group differences in time spent in the OA of the EPM (Fig. 5B) or percent habituation of the startle (Fig. 5C). Analysis of freezing behavior across the three days of testing revealed a significant Session x Dose interaction [F_(2, 22)_ = 4.771, p = 0.0190]. Post-hoc analyses indicated that resilient rats treated with intra-BLA MTEP (10μg/side) displayed a higher amount of freezing relative to vehicle-treated Resilient rats on Day 1 (p < 0.01). On Days 2 and 3 of re-exposure, no MTEP was delivered and freezing declined, with a significant difference between Day 1 and Day 3 in the group that had received MTEP on Day 1 (p < 0.05).

**Figure 5.**
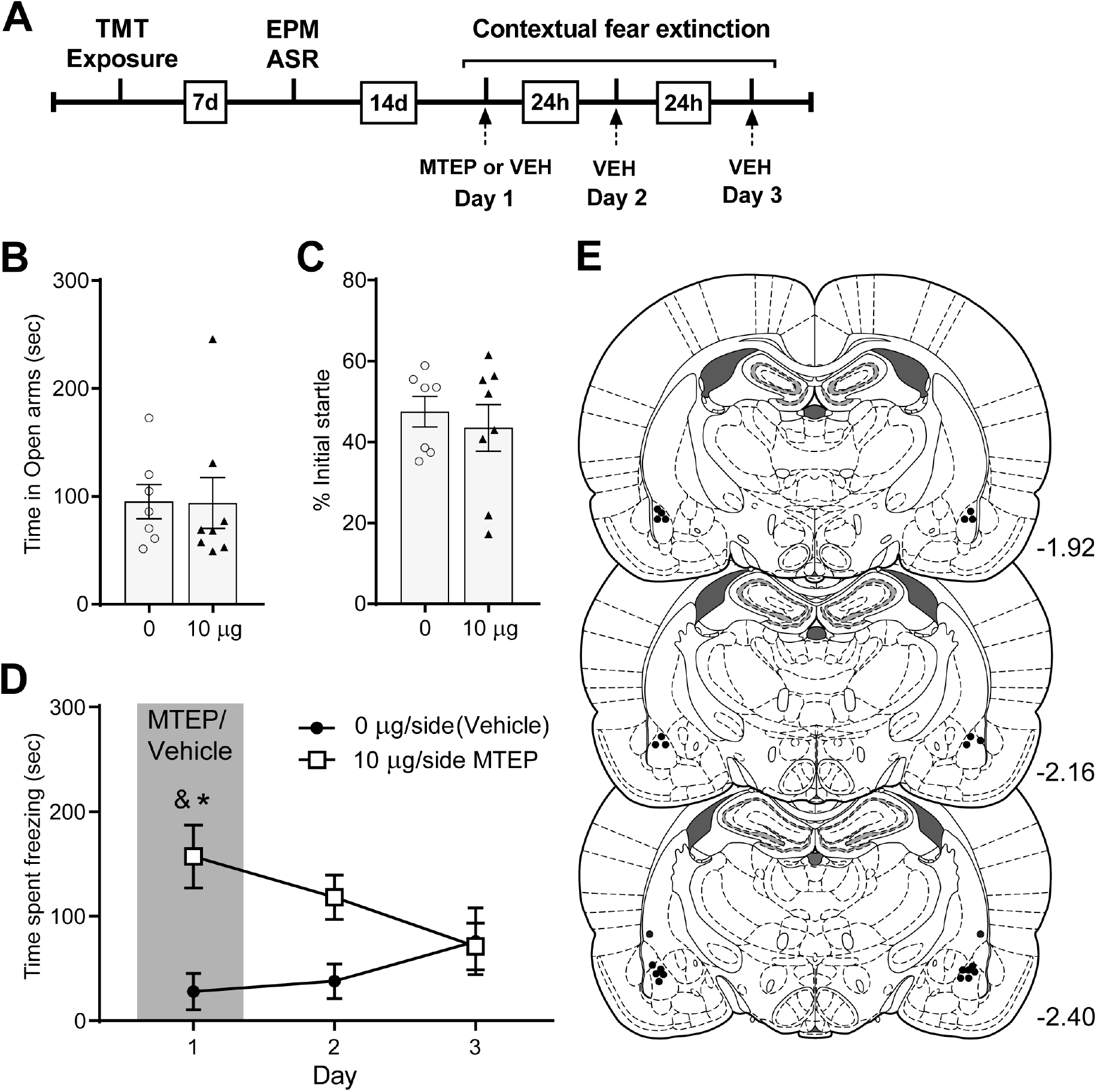
mGlu5 receptors in the BLA mediate contextual fear. **A.** Timeline of Experiment 2. **B.** Time spent in the open arms of the EPM and percent initial startle response **(C)** did not differ between rats later assigned to receive vehicle or MTEP. **D.** On the day of administration (Day 1) MTEP increased freezing in the TMT environment, which declined by extinction Day 3. * = p<0.01 vs. Vehicle; & = p<0.05 vs. Day 3. Black circles indicate the tip of the injectors within the BLA; one Vehicle-treated rat had placement that was dorsal to the BLA, but data was not excluded. Coronal outlines are based on the rat brain atlas (Paxinos and Watson, 2007).

## Discussion

We have previously reported greater expression of mGlu5 mRNA in the amygdala and mPFC and higher CB1 receptor mRNA in the amygdala of rats shown to be Resilient to long-term fear/anxiety following a single exposure to predator scent stress (TMT), relative to both unstressed Controls and stress-Susceptible rats (Schwendt et al. 2018). The present study extends these findings in three fundamental ways. First, by examining the sub-regional and cell-type-specific expression of mGlu5 and CB1R mRNAs using multiplex fluorescent in situ hybridization (FISH) in both brain regions; second, by assessing the possible effect of re-exposure to TMT context on receptor mRNA levels, and third, by evaluating the effects of BLA-specific mGlu5 inhibition on conditioned fear responses in Resilient rats

In the mPFC, the FISH analysis revealed elevated mGlu5 mRNA in glutamatergic PNs in both IL and PL regions in Resilient rats relative to unstressed Controls, independent of re-exposure to the TMT context. We also found striking increases in BLA mGlu5 mRNA localized in glutamatergic PNs in Resilient rats. Increased BLA CB1 mRNA was found in the low expressing population of vGlut1-containing PNs, and no difference in expression in non-GABAergic (high expressing) populations. There were no differences in mGlu5 or CB1 expression in the CeA, nor did context re-exposure affect expression in any brain region examined. We chose to use the Control condition as a comparison group for mGlu5 and CB1 mRNA assessment because our prior work found no differences in expression of these genes between Controls and Susceptible rats. It is possible that here we would have found different results if we compared sub-region, cell-type-specific expression between Resilient and Susceptible rats. However, our goal was to characterize these adaptations in stress-Resilient rats in order to understand that particular phenotype, not to investigate the neurobiology of Susceptibility. Finally, the necessity of mGlu5 signaling in the BLA for mediating extinction of conditioned fear in Resilient rats was demonstrated by an increase in freezing on the day of intra-BLA MTEP administration, which extinguished over subsequent days.

Increased mGlu5 expression in the IL as a feature of the Resilient phenotype in this rodent model is consistent with preclinical findings demonstrating a critical role for mGlu5 in the consolidation of extinction memory. During extinction recall, activation of mGlu5 receptors promotes increased intrinsic excitability of IL neurons by causing a prolonged reduction in the slow afterhyperpolarization currents (Fontanez-Nuin et al. 2011). In addition, mGlu5 is involved in extinction-evoked LTP at IL synapses by supporting an increased AMPA/NMDA ratio, AMPAR rectification index, and CP-AMPAR conductance (Sepulveda-Orengo et al. 2013). These mGlu5 mediated changes serve to enhance the responsiveness of IL neurons to afferent signaling providing new information regarding the conditioned stimulus, subsequently supporting the burst-firing of IL neurons projecting towards inhibitory amygdalar nuclei and effectively shunting pro ‘fear’ signals arriving from the PL (Vogel et al. 2016).

Stimulation of ascending glutamatergic BLA “extinction” neurons is necessary for excitation of the IL and subsequent extinction (Herry et al. 2008; Pape and Pare 2010). During fear-extinction, augmented excitability of BLA neurons projecting to the IL results in glutamate release and burst-firing of IL PNs which in turn promotes long-lasting potentiation of synapses at downstream inhibitory amygdalar subnuclei (Quirk et al. 2003; Fontanez-Nuin et al. 2011) Increased excitability of these newly strengthened circuits contributes to a shunting of excitatory ‘fear’ signals arriving from the PL, effectively countering PL mediated activation of the CeA and associated physiological fear response (Thompson et al. 2010). Despite observations of mGlu5 mediated LTP in the BLA (Rodrigues et al. 2002; Chen et al. 2017; Rahman et al. 2017) whether these receptors are involved in extinction-evoked excitability of IL targeting BLA neurons is not clear. Using a model of spontaneous recovery of footshock-conditioned fear, Mao et al. (2013) found that both systemic (2.5 and 5 mg/kg) and intra-BLA (10 μg) infusion of mGlu5 antagonist MTEP prior to extinction training produced deficits in extinction recall which were linked to increased GluR1 surface expression and AMPA/NMDA ratios. These findings support a more traditional “dual” role for mGlu5 in promoting LTD at BLA synapses during extinction learning and LTP during fear conditioning (Ayala et al. 2009; Chen et al. 2017). Nonetheless, the fact remains that mGlu5 receptor signaling dynamics in the BLA during extinction are complex and only marginally understood (Niswender and Conn 2010; Rook et al. 2015). Recent studies have demonstrated that mGlu5 and a second Group I mGluR, mGlu1, receptor function in the BLA is contingent upon stimulus intensity, measured as glutamate concentrations. This phenomenon, known as “metaplasticity”, critically influences mGlu1/mGlu5 signaling transduction and dependence of LTP and LTD on NMDA receptor function in the BLA (Abraham 2008; Rudy and Matus-Amat 2009; Chen et al. 2017)

Further illustrating multimodal mGlu5 function in the BLA, a recent study found that the same populations of receptors modulate both associative fear learning and indiscriminate anxiety (Rahman et al. 2017). Thus, stimulation of mGlu5 at levels below the threshold for triggering LTP results in anxiety-like behavior in rodents independent of learning. Indeed, intra-BLA infusion of mGlu5 antagonist reduces anxiety-like behaviors in rodents and systemic administration of mGlu5 antagonist is anxiolytic in humans (Porter et al. 2005; De La Mora et al. 2006). Furthermore, stimulation of mGlu5 paired with presentation of cue-specific stimuli is sufficient to produce above-threshold activation triggering LTP induction. These findings are in line with recent electrophysiological characterization of BLA LTP, which indicates that mGlu5 “priming” of BLA neurons through mobilization of intracellular calcium and increased membrane excitability is necessary for NMDA mediated calcium influx to reach concentrations required for LTP (Chen et al. 2017). Thus, our finding that intra-BLA administration of 10ug MTEP prior to context re-exposure served to increase freezing in Resilient rats suggests that mGlu5 blockade prevents initiation of synaptic plasticity mechanisms involved in acquisition and consolidation of extinction. From these findings one might propose that the increased mGlu5 mRNA tone in Resilient rats serves to augment mGlu5-mediated plasticity involved in fear learning, providing them with a mechanism for “rapid” and enduring extinction, which occurs immediately in the absence of mGlu5 blockade.

Stimulation of mGlu5 receptors (among other things) triggers the de novo synthesis of the CB1 agonist 2-Arachidonoylglycerol (2-AG), initiating short-term plasticity through producing rapid reductions in synaptic glutamate release (Castillo et al. 2012). Following the identification of distinct ‘fear’ and ‘extinction’ PNs in the BLA, a handful of comprehensive studies have set out to characterize the role of CB1R containing CCK+ basket cells in bidirectionally regulating the excitability of these oppositional cell groups (Herry et al. 2008; Tan et al. 2011; Vogel et al. 2016). Although there is similar connectivity between the CCK^+^ basket cells and both IL and PL projecting populations of BLA PNs, terminals neighboring postsynaptic regions of the IL projecting “extinction” population have been shown to produce stronger DSI, which is thought to be a result of upregulated diacylglycerol lipase alpha (DAGLα – a precursor to 2-AG synthesis) in this PN population (Wilson and Nicoll 2001; Vogel et al. 2016). To our surprise, we found no differences in CB1R mRNA expression in the GABAergic population of high CB1-expressing cells in the BLA of Resilient rats relative to Controls, despite Resilient rats consistently exhibit rapid contextual extinction (Schwendt et al., 2018; Fig. 5). We did, however, find increased CB1 mRNA in the low-expressing population of glutamatergic BLA PNs, implicating an augmented expression of CB1R at axonal terminals at brain regions receiving BLA projections. It is possible that the increase in PN CB1 mRNA expression is localized in PN targeting cells, which could potentially result in reduced release of glutamate across fear promoting BLA to PL pathways. Alternatively, it was recently demonstrated that increased CB1 mRNA in BLA PNs with terminals targeting dopamine receptor type 2 containing MSNs in the nucleus accumbens prevented long-term depressive behaviors following social defeat in mice, essentially promoting stress ‘resiliency’ (Shen et al. 2019).

### Conclusions

Here we demonstrate that rats resilient to long-term anxiety/fear produced by a single predatory scent stress exposure display upregulated mGlu5 mRNA in glutamatergic neurons of the BLA, PL and IL, and upregulated CB1 in glutamatergic neurons of the BLA. Future work will seek to investigate the interaction between CB1 and mGlu5 receptors in the BLA and in BLA afferents.

## Acknowledgments

The authors thank Jason Dee, Stephen Beaudin-Curley and Doug Smith, for their assistance with image acquisition and analysis. The fluorescent microscopy images were acquired using Olympus DSU (DU-DBIX) spinning disk confocal microscope located at the Cell & Tissue Analysis Core (CTAC) facility. This facility is funded through user fees and the generous financial support of the McKnight Brain Institute at the University of Florida. This research was supported by the following grants: the subcontract 8738sc and 9250sc from the Institute on Molecular Neuroscience (awarded to LAK). Award Number: W81XWH-12-2-0048. The U.S. Army Medical Research Acquisition Activity, 820 Chandler Street, Fort Detrick, MD 21702-5014 is the awarding and administering acquisition office. The content of the information does not necessarily reflect the position or the policy of the Government, and no official endorsement should be inferred; and by the pilot grant from the Center for OCD, Anxiety, and Related Disorders (COARD) at the University of Florida (awarded to MS).

## Author contributions

JS, performed all behavioral and in situ hybridization experiments, wrote the first draft of the manuscript, compiled, and edited the figures, and edited successive drafts of the manuscript. LW, assisted with behavioral experiments and processed brain tissue for the histological analysis. LAK and MS co-designed study, co-directed the research, oversaw all aspects of data analysis and edited successive drafts of the manuscript. MS, prepared the final version of the manuscript.

## Compliance with ethical standards

Conflict of interest All authors declare no conflict of interest.

## References

Abraham WC (2008) Metaplasticity: Tuning synapses and networks for plasticity. Nat. Rev. Neurosci. 9:387–399

Ayala JE, Chen Y, Banko JL, et al (2009) MGluR5 Positive Allosteric Modulators Facilitate both Hippocampal LTP and LTD and Enhance Spatial Learning. Neuropsychopharmacology 34:2057–2071. https://doi.org/10.1038/npp.2009.30

Bahar Halpern K, Itzkovitz S (2016) Single molecule approaches for quantifying transcription and degradation rates in intact mammalian tissues. Methods 98:134–142. https://doi.org/10.1016/j.ymeth.2015.11.015

Blechert J, Michael T, Vriends N, et al (2007) Fear conditioning in posttraumatic stress disorder: evidence for delayed extinction of autonomic, experiential, and behavioural responses. Behav Res Ther 45:2019–33. https://doi.org/10.1016/j.brat.2007.02.012

Bloodgood DW, Sugam JA, Holmes A, Kash TL (2018) Fear extinction requires infralimbic cortex projections to the basolateral amygdala. Transl Psychiatry 8:60. https://doi.org/10.1038/s41398-018-0106-x

Bouton ME, Westbrook RF, Corcoran KA, Maren S (2006) Contextual and Temporal Modulation of Extinction: Behavioral and Biological Mechanisms. Biol. Psychiatry 60:352–360

Breslau N, Davis GC, Andreski P, Peterson E (1991) Traumatic Events and Posttraumatic Stress Disorder in an Urban Population of Young Adults. Arch Gen Psychiatry 48:216–222. https://doi.org/10.1001/archpsyc.1991.01810270028003

Bukalo O, Pinard CR, Silverstein S, et al (2015) Prefrontal inputs to the amygdala instruct fear extinction memory formation. Sci Adv 1:e1500251. https://doi.org/10.1126/sciadv.1500251

Caballero JP, Scarpa GB, Remage-Healey L, Moorman DE (2019) Differential effects of dorsal and ventral medial prefrontal cortex inactivation during natural reward seeking, extinction, and cue-induced reinstatement. eNeuro 6:. https://doi.org/10.1523/ENEURO.0296-19.2019

Castillo PE, Younts TJ, Chávez AE, Hashimotodani Y (2012) Endocannabinoid Signaling and Synaptic Function. Neuron 76:70–81

Chen A, Hu WW, Jiang XL, et al (2017) Molecular mechanisms of group I metabotropic glutamate receptor mediated LTP and LTD in basolateral amygdala in vitro. Psychopharmacology (Berl) 234:681–694. https://doi.org/10.1007/s00213-016-4503-7

Cho JH, Deisseroth K, Bolshakov VY (2013) Synaptic encoding of fear extinction in mPFC-amygdala circuits. Neuron 80:1491–1507. https://doi.org/10.1016/j.neuron.2013.09.025

Ciocchi S, Herry C, Grenier F, et al (2010) Encoding of conditioned fear in central amygdala inhibitory circuits. Nature 468:277–282. https://doi.org/10.1038/nature09559

Day HEW, Masini C V., Campeau S (2004) The pattern of brain c-fos mRNA induced by a component of fox odor, 2,5-dihydro-2,4,5-Trimethylthiazoline (TMT), in rats, suggests both systemic and processive stress characteristics. Brain Res 1025:139–151. https://doi.org/10.1016/j.brainres.2004.07.079

De La Mora MP, Lara-García D, Jacobsen KX, et al (2006) Anxiolytic-like effects of the selective metabotropic glutamate receptor 5 antagonist MPEP after its intra-amygdaloid microinjection in three different non-conditioned rat models of anxiety. Eur J Neurosci 23:2749–2759. https://doi.org/10.1111/j.1460-9568.2006.04798.x

Duvarci S, Pare D (2014) Amygdala microcircuits controlling learned fear. Neuron 82:966–980

Ehrlich I, Humeau Y, Grenier F, et al (2009) Amygdala Inhibitory Circuits and the Control of Fear Memory. Neuron 62:757–771

Esterlis I, Holmes SE, Sharma P, et al (2018) Metabotropic Glutamatergic Receptor 5 and Stress Disorders: Knowledge Gained From Receptor Imaging Studies. Biol Psychiatry 84:95–105. https://doi.org/10.1016/j.biopsych.2017.08.025

Ferraguti F, Shigemoto R (2006) Metabotropic glutamate receptors. Cell Tissue Res. 326:483–504

Fitzgerald ML, Mackie K, Pickel VM (2019) Ultrastructural localization of cannabinoid CB1 and mGluR5 receptors in the prefrontal cortex and amygdala. J Comp Neurol 527:2730–2741. https://doi.org/10.1002/cne.24704

Fontanez-Nuin DE, Santini E, Quirk GJ, Porter JT (2011) Memory for fear extinction requires mGluR5-mediated activation of infralimbic neurons. Cereb Cortex 21:727–735. https://doi.org/10.1093/cercor/bhq147

Gale GD, Anagnostaras SG, Godsil BP, et al (2004) Role of the Basolateral Amygdala in the Storage of Fear Memories across the Adult Lifetime of Rats. J Neurosci 24:3810–3815. https://doi.org/10.1523/JNEUROSCI.4100-03.2004

Guthrie RM, Bryant RA (2006) Extinction learning before trauma and subsequent posttraumatic stress. Psychosom Med 68:307–311. https://doi.org/10.1097/01.psy.0000208629.67653.cc

Herry C, Ciocchi S, Senn V, et al (2008) Switching on and off fear by distinct neuronal circuits. Nature 454:600–606. https://doi.org/10.1038/nature07166

Heuss C, Scanziani M, Gähwiler BH, Gerber U (1999) G-protein-independent signaling mediated by metabotropic glutamate receptors. Nat Neurosci 2:1070–1077. https://doi.org/10.1038/15996

Holmes SE, Girgenti MJ, Davis MT, et al (2017) Altered metabotropic glutamate receptor 5 markers in PTSD: In vivo and postmortem evidence. Proc Natl Acad Sci 114:8390–8395. https://doi.org/10.1073/pnas.1701749114

Jasnow AM, Ressler KJ, Hammack SE, et al (2009) Distinct subtypes of cholecystokinin (CCK)-containing interneurons of the basolateral amygdala identified using a CCK promoter-specific lentivirus. J Neurophysiol 101:1494–1506. https://doi.org/10.1152/jn.91149.2008

Kano M, Ohno-Shosaku T, Hashimotodani Y, et al (2009) Endocannabinoid-mediated control of synaptic transmission. Physiol. Rev. 89:309–380

Katona I, Rancz EA, Acsády L, et al (2001) Distribution of CB1 cannabinoid receptors in the amygdala and their role in the control of GABAergic transmission. J Neurosci 21:9506–9518. https://doi.org/10.1523/jneurosci.21-23-09506.2001

Kheirbek MA, Drew LJ, Burghardt NS, et al (2013) Differential control of learning and anxiety along the dorsoventral axis of the dentate gyrus. Neuron 77:955–968. https://doi.org/10.1016/j.neuron.2012.12.038

Kim W Bin, Cho JH (2017) Encoding of Discriminative Fear Memory by Input-Specific LTP in the Amygdala. Neuron 95:1129–1146.e5. https://doi.org/10.1016/j.neuron.2017.08.004

Koenigs M, Grafman J (2009) Posttraumatic stress disorder: The role of medial prefrontal cortex and amygdala. Neuroscientist 15:540–548

Krabbe S, Gründemann J, Lüthi A (2018) Amygdala Inhibitory Circuits Regulate Associative Fear Conditioning. Biol. Psychiatry 83:800–809

Laricchiuta D, Saba L, De Bartolo P, et al (2016) Maintenance of aversive memories shown by fear extinction-impaired phenotypes is associated with increased activity in the amygdaloid-prefrontal circuit. Sci Rep 6:1–13. https://doi.org/10.1038/srep21205

Mao S-C, Chang C-H, Wu C-C, et al (2013) Inhibition of Spontaneous Recovery of Fear by mGluR5 after Prolonged Extinction Training. PLoS One 8:e59580. https://doi.org/10.1371/journal.pone.0059580

Marcus DJ, Bedse G, Gaulden AD, et al (2020) Endocannabinoid Signaling Collapse Mediates Stress-Induced Amygdalo-Cortical Strengthening. Neuron 105:1062–1076.e6. https://doi.org/10.1016/j.neuron.2019.12.024

Marsicano G, Lutz B (1999) Expression of the cannabinoid receptor CB1 in distinct neuronal subpopulations in the adult mouse forebrain. Eur J Neurosci 11:4213–4225. https://doi.org/10.1046/j.1460-9568.1999.00847.x

McGarry LM, Carter AG (2017) Prefrontal Cortex Drives Distinct Projection Neurons in the Basolateral Amygdala. Cell Rep 21:1426–1433. https://doi.org/10.1016/j.celrep.2017.10.046

Muly EC, Maddox M, Smith Y (2003) Distribution of mGluR1α and mGluR5 Immunolabeling in Primate Prefrontal Cortex. J Comp Neurol 467:521–535. https://doi.org/10.1002/cne.10937

Neumeister A, Normandin MD, Pietrzak RH, et al (2013) Elevated brain cannabinoid CB 1 receptor availability in post-traumatic stress disorder: A positron emission tomography study. Mol Psychiatry 18:1034–1040. https://doi.org/10.1038/mp.2013.61

Niswender CM, Conn PJ (2010) Metabotropic Glutamate Receptors: Physiology, Pharmacology, and Disease. https://doi.org/10.1146/annurev.pharmtox.011008.145533

Orr SP, Metzger LJ, Lasko NB, et al (2000) De novo conditioning in trauma-exposed individuals with and without posttraumatic stress disorder. J Abnorm Psychol 109:290–298. https://doi.org/10.1037/0021-843X.109.2.290

Pape HC, Pare D (2010) Plastic synaptic networks of the amygdala for the acquisition, expression, and extinction of conditioned fear. Physiol. Rev. 90:419–463

Paxinos G, Watson C (2007) The Rat Brain in Stereotaxic Coordinates Sixth Edition. Boston, MA

Perkonigg A, Kessler RC, Storz S, Wittchen HU (2000) Traumatic events and posttraumatic stress disorder in the community: Prevalence, risk factors and comorbidity. Acta Psychiatr Scand 101:46–59. https://doi.org/10.1034/j.1600-0447.2000.101001046.x

Pietrzak RH, Huang Y, Corsi-Travali S, et al (2014) Cannabinoid type 1 receptor availability in the amygdala mediates threat processing in trauma survivors. Neuropsychopharmacology 39:2519–2528. https://doi.org/10.1038/npp.2014.110

Porter RHP, Jaeschke G, Spooren W, et al (2005) Fenobam: A clinically validated nonbenzodiazepine anxiolytic is a potent, selective, and noncompetitive mGlu5 receptor antagonist with inverse agonist activity. J Pharmacol Exp Ther 315:711–721. https://doi.org/10.1124/jpet.105.089839

Quirk GJ, Likhtik E, Pelletier JG, Paré D (2003) Stimulation of medial prefrontal cortex decreases the responsiveness of central amygdala output neurons. J Neurosci 23:8800–8807. https://doi.org/10.1523/jneurosci.23-25-08800.2003

Rahman MM, Kedia S, Fernandes G, Chattarji S (2017) Activation of the same mGluR5 receptors in the amygdala causes divergent effects on specific versus indiscriminate fear. Elife 6:. https://doi.org/10.7554/eLife.25665

Rodrigues SM, Bauer EP, Farb CR, et al (2002) The group I metabotropic glutamate receptor mGluR5 is required for fear memory formation and long-term potentiation in the lateral amygdala. J Neurosci 22:5219–29. https://doi.org/10.1523/JNEUROSCI.22-12-05219.2002

Rook JM, Xiang Z, Lv X, et al (2015) Biased mGlu5-Positive Allosteric Modulators Provide InVivo Efficacy without Potentiating mGlu5 Modulation of NMDAR Currents. Neuron 86:1029–1040. https://doi.org/10.1016/j.neuron.2015.03.063

Rudy JW, Matus-Amat P (2009) DHPG activation of group 1 mGluRs in BLA enhances fear conditioning. Learn Mem 16:421–425. https://doi.org/10.1101/lm.1444909

Sah P, Faber ESL, De Armentia ML, Power J (2003) The amygdaloid complex: Anatomy and physiology. Physiol. Rev. 83:803–834

Sareen J (2014) Posttraumatic stress disorder in adults: Impact, comorbidity, risk factors, and treatment. Can. J. Psychiatry 59:460–467

Schneider CA, Rasband WS, Eliceiri KW (2012) NIH Image to ImageJ: 25 years of image analysis. Nat. Methods 9:671–675

Schwendt M, Shallcross J, Hadad NA, et al (2018) A novel rat model of comorbid PTSD and addiction reveals intersections between stress susceptibility and enhanced cocaine seeking with a role for mGlu5 receptors. Transl Psychiatry 8:209. https://doi.org/10.1038/s41398-018-0265-9

Sepulveda-Orengo MT, Lopez A V., Soler-Cedeño O, Porter JT (2013) Fear extinction induces mGluR5-mediated synaptic and intrinsic plasticity in infralimbic neurons. J Neurosci 33:7184–7193. https://doi.org/10.1523/JNEUROSCI.5198-12.2013

Shallcross J, Hámor P, Bechard AR, et al (2019) The Divergent Effects of CDPPB and Cannabidiol on Fear Extinction and Anxiety in a Predator Scent Stress Model of PTSD in Rats. Front Behav Neurosci 13:. https://doi.org/10.3389/fnbeh.2019.00091

Shen CJ, Zheng D, Li KX, et al (2019) Cannabinoid CB1 receptors in the amygdalar cholecystokinin glutamatergic afferents to nucleus accumbens modulate depressive-like behavior. Nat Med 25:337–349. https://doi.org/10.1038/s41591-018-0299-9

Sierra-Mercado D, Padilla-Coreano N, Quirk GJ (2011) Dissociable Roles of Prelimbic and Infralimbic Cortices, Ventral Hippocampus, and Basolateral Amygdala in the Expression and Extinction of Conditioned Fear. Neuropsychopharmacology 36:529–538. https://doi.org/10.1038/npp.2010.184

Sotres-Bayon F, Cain CK, LeDoux JE (2006) Brain Mechanisms of Fear Extinction: Historical Perspectives on the Contribution of Prefrontal Cortex. Biol. Psychiatry 60:329–336

Tan H, Lauzon NM, Bishop SF, et al (2011) Cannabinoid transmission in the basolateral amygdala modulates fear memory formation via functional inputs to the prelimbic cortex. J Neurosci 31:5300–5312. https://doi.org/10.1523/JNEUROSCI.4718-10.2011

Thompson BM, Baratta M V., Biedenkapp JC, et al (2010) Activation of the infralimbic cortex in a fear context enhances extinction learning. Learn Mem 17:591–599. https://doi.org/10.1101/lm.1920810

Vogel E, Krabbe S, Gründemann J, et al (2016) Projection-Specific Dynamic Regulation of Inhibition in Amygdala Micro-Circuits. Neuron 91:644–651. https://doi.org/10.1016/j.neuron.2016.06.036

Vouimba RM, Maroun M (2011) Learning-induced changes in mpfc-bla connections after fear conditioning, extinction, and reinstatement of fear. Neuropsychopharmacology 36:2276–2285. https://doi.org/10.1038/npp.2011.115

Wang F, Flanagan J, Su N, et al (2012) RNAscope: A novel in situ RNA analysis platform for formalin-fixed, paraffin-embedded tissues. J Mol Diagnostics 14:22–29. https://doi.org/10.1016/j.jmoldx.2011.08.002

Wang H, Zhuo M (2012) Group I Metabotropic Glutamate Receptor-Mediated Gene Transcription and Implications for Synaptic Plasticity and Diseases. Front Pharmacol 3:1–8. https://doi.org/10.3389/fphar.2012.00189

Wilson RI, Nicoll RA (2001) Endogenous cannabinoids mediate retrograde signalling at hippocampal synapses. Nature 410:588–592. https://doi.org/10.1038/35069076

Yoshida T, Uchigashima M, Yamasaki M, et al (2011) Unique inhibitory synapse with particularly rich endocannabinoid signaling machinery on pyramidal neurons in basal amygdaloid nucleus. Proc Natl Acad Sci U S A 108:3059–3064. https://doi.org/10.1073/pnas.1012875108

Zhu PJ, Lovinger DM (2005) Retrograde endocannabinoid signaling in a postsynaptic neuron/synaptic bouton preparation from basolateral amygdala. J Neurosci 25:6199–6207. https://doi.org/10.1523/JNEUROSCI.1148-05.2005

Zou S, Kumar U (2018) Cannabinoid Receptors and the Endocannabinoid System: Signaling and Function in the Central Nervous System. Int J Mol Sci 19:833. https://doi.org/10.3390/ijms19030833

